# A central role for Numb/Nbl in multiple Shh-mediated axon repulsion processes

**DOI:** 10.1101/2024.04.29.591732

**Authors:** Tiphaine Dolique, Sarah Baudet, Frederic Charron, Julien Ferent

## Abstract

Sonic hedgehog (Shh) is an axon guidance molecule that can act as either a chemorepellent or a chemoattractant, depending on the neuron type and their developmental stage. In the developing spinal cord, Shh initially attracts commissural axons to the floor plate and later induces their repulsion after they cross the midline. In the developing visual system, Shh repels ipsilateral retinal ganglion cell (iRGC) axons at the optic chiasm. Although Shh requires the endocytic adaptor Numb for attraction of commissural neurons, the molecular mechanisms underlying Shh dual function in attraction and repulsion are still unclear. In this study, we investigate whether Numb also regulates repulsive axon guidance. We show that Numb is essential for two Shh-mediated repulsion processes: iRGC axon repulsion at the optic chiasm and antero-posterior commissural axon repulsion in the spinal cord. Therefore, Numb is required for Shh-mediated attraction and repulsion. These results position Numb as a central player in the non-canonical Shh signalling pathway mediating axon repulsion.

**Summary statement:** Here, we show that Numb is required for Shh-mediated midline repulsion of ipsilateral retinal ganglion cell axons and post-crossing commissural axons of the spinal cord.

## INTRODUCTION

During embryogenesis, growing axons are guided to their targets by molecular guidance cues. Sensing these cues, whether they are repulsive or attractive, is essential for correct axon guidance and circuit formation (Tessier-Lavigne and Goodman, 1996). Among the many guidance cues, Sonic Hedgehog (Shh) is a critical molecule that can act as a chemorepellent or chemoattractant, depending on the neuron type and developmental stage. In the neural tube, Shh is initially responsible for the attraction of commissural axons to the floor plate, through the activation of Smoothened (Smo) and Src family kinases (SFKs) (Charron et al., 2003; Okada et al., 2006; Yam et al., 2009). After they have crossed the midline, Shh repels commissural axons along the antero-posterior axis (Yam et al., 2012).

In the developing chick visual system, a species devoid of binocular vision, Shh is a chemorepulsive cue for retinal ganglion cell (RGC) axons (Trousse et al., 2001). In species with binocular vision, RGCs can either project to the same side of the brain (ipsilateral, iRGC) or to the opposite side (contralateral, cRGC). The segregation of these two types of RGCs occurs at the level of the optic chiasm and is controlled by the repulsion of iRGCs by Ephrins (through their Eph receptors) and by Shh (through its receptor Boc) (Fabre et al., 2010; Herrera et al., 2024; Herrera et al., 2018; Murcia-Belmonte and Erskine, 2019; Petros et al., 2009; Williams et al., 2003). Shh is produced by cRGCs in the retina and is then transported anterogradely along the axon (Peng et al., 2018). Shh accumulates at the optic chiasm and induces repulsion of iRGCs via an axon-axon interaction mechanism.

Receptors are targeted to specific sites on the cell membrane through endocytosis and/or exocytosis, a mechanism essential for axon guidance (Tojima and Kamiguchi, 2015; Winckler and Mellman, 2010; Yap and Winckler, 2012; Yap and Winckler, 2015). We have previously shown that Shh-mediated Boc endocytosis is required for the attraction of commissural axons in the developing spinal cord (Ferent et al., 2019). Both Shh attraction and Boc internalization are dependent on the endocytic membrane-associated protein Numb. However, it is unknown whether the function of Numb in Shh signaling is conserved in other non-canonical Shh responses such as axonal repulsion.

Therefore, we investigated whether Numb is essential for axon repulsion in two different systems where Shh acts as a repulsive cue: the segregation of ipsilaterally-projecting RGC axons at the optic chiasm and the antero-posterior repulsion of post-crossing commissural axons after they have crossed the neural tube midline.

## RESULTS

### Numb and Nbl are expressed in iRGCs

Numb and its homolog Numb-like (Nbl) are expressed broadly the developing nervous system (Zhong et al., 1997) and can be functionally redundant (Petersen et al., 2002). We examined the expression of Numb and Nbl in the retina using an antibody that recognizes both proteins (Kechad et al., 2012). Numb/Nbl expression has been described in the RGC layer (Kechad et al., 2012), but it is unknown whether it is expressed by iRGCs. To specifically label iRGCs, we used a Cre recombinase under the control of the Slc6a4 promoter (serotonin transporter; Sert-Cre) to activate the expression of the tdTomato reporter gene (Figure 1A) (Koch et al., 2013; Peng et al., 2018). In the retina of postnatal day 0 (P0) mouse, tdTomato^+^ cells are restricted to the dorsolateral region, where iRGCs are located (Figure 1B, B’). Numb/Nbl expression appears high throughout the retina, especially in the RGC layer (Figure 1B, and we detected a strong colocalization of Numb/Nbl with tdTomato^+^ cells (Figure 1B’), demonstrating robust expression of Numb/Nbl by iRGCs. We observed that Numb/Nbl signal is also very high in the optic nerve and at the optic chiasm (Figure 1C, D). In addition to labelling cell bodies, the tdTomato signal allows visualization of iRGC axons along the entire optic pathway. Colocalization of Numb/Nbl and tdTomato is also high in axons in the optic nerve and chiasm (Figure 1C’, D). Axonal localization of Numb/Nbl has also been observed in other types of neurons, such as hippocampal neurons (Nishimura et al., 2003) or spinal commissural neurons (Ferent et al., 2019), suggesting that Numb/Nbl may also play a role at a distance from the cell bodies of iRGCs.

**Figure 1:**
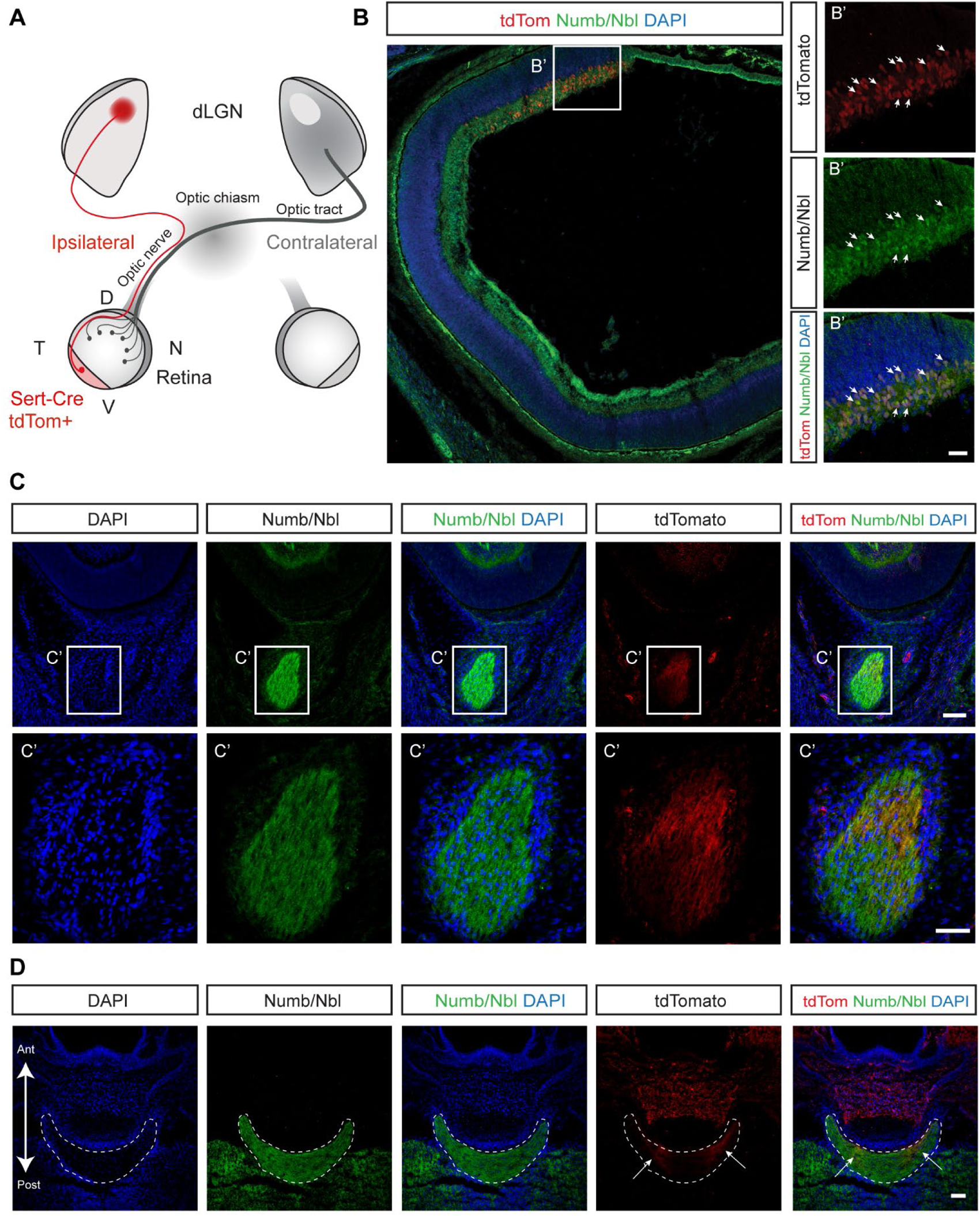
Numb/Nbl is expressed by iRGCs. (A) Schematic showing the organization of RGC projections from the retina to the dLGN and the distribution of tdTomato expressed under the Sert-Cre driver. (B) Mouse P0 Sert-Cre; Rosa26-tdTomato retina sections immunostained for Numb/Nbl. (B’) Enlargement of the retinal VT region where tdTomato^+^ cells are found. Neurons positive for both tdTomato and Numb/Nbl are indicated by white arrows. (C) Optic nerve sections immunostained for Numb/Nbl. (C’) Enlargement of the optic nerve section where tdTomato^+^ axons are detected, together with Numb/Nbl staining. (D) Optic chiasm sections (dotted line) immunostained for Numb/Nbl. The arrows show Sert-Cre; Rosa26-tdTomato^+^ ipsilateral projections from the retina. Scale bars: 100 μm in B; 25 μm in B’; 100 μm in C and C’; 100 μm in D.

### Numb/Nbl iRGC conditional knock-out does not affect eye morphology and iRGC specification

When *Numb* and *Nbl* are deleted in the early retina, eye development is severely disrupted (Kechad et al., 2012). Mutant eyes are hypoplastic and the number of most retinal cell types is reduced (Kechad et al., 2012). To investigate the role of Numb and Nbl specifically in iRGC and avoid the retinal structural defects caused by early deletion of *Numb* and *Nbl*, we used the Sert-Cre mouse line to inactivate conditional alleles of *Numb* and *Nbl* (Wilson et al., 2007). We refer to the resulting conditional mutants as Sert-Cre cDKO. The structure of the eyes of P0 Sert-Cre cKOs was analyzed in comparison to control littermate animals. Measurements of several parameters including eye diameter, eye area, retinal area, optic nerve and optic tract area showed no differences between Sert-Cre cDKO and controls (Figure 2A-F). Thus, conditional knockout of *Numb* and *Nbl* in iRGCs does not affect the gross anatomy and morphological measurements of the eye and its projection.

**Figure 2:**
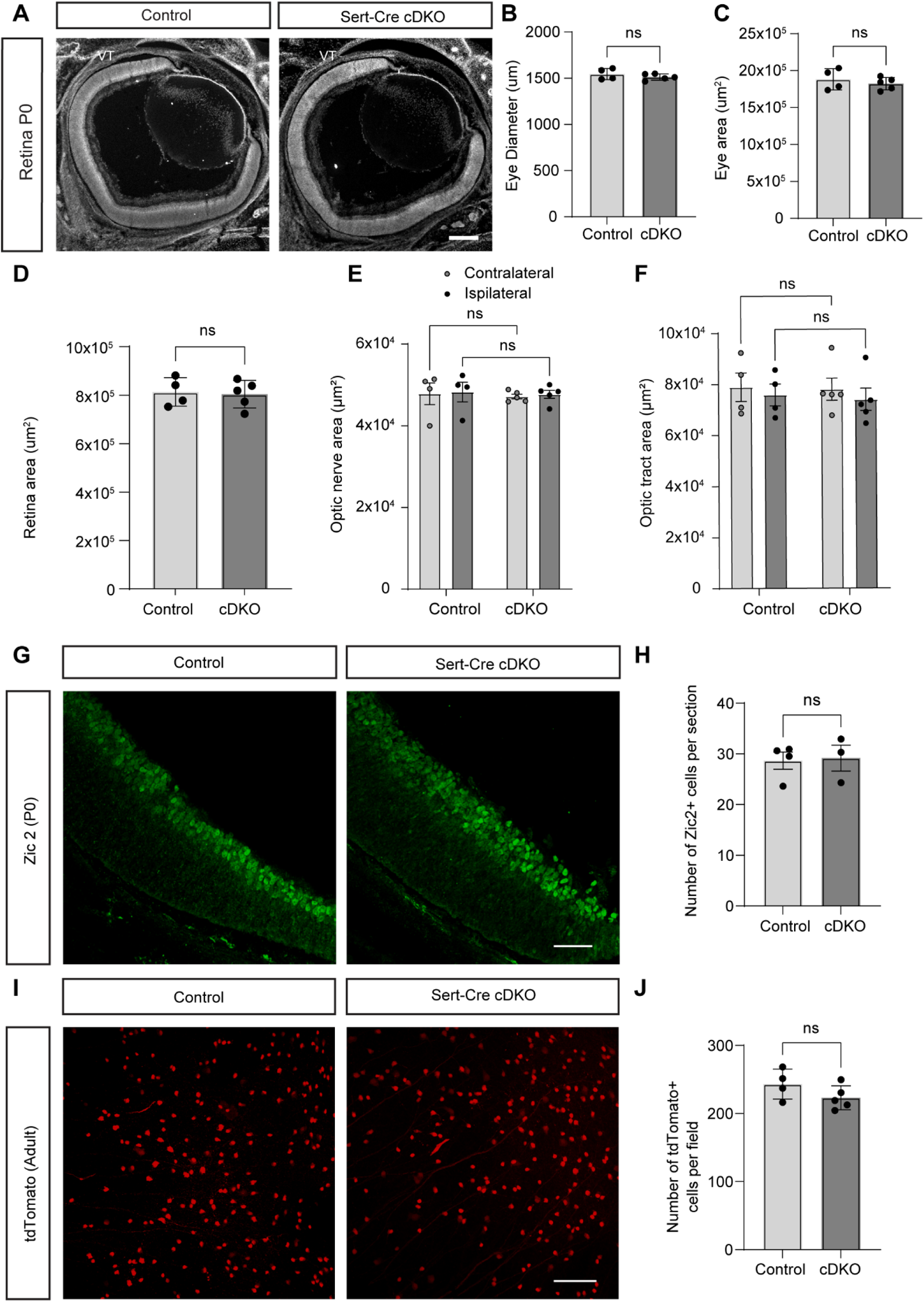
Numb/Nbl cDKO in iRGCs does not affect eye morphology and iRGC specification. (A) Retina sections at P0 from control and Sert-Cre cDKO animals. The cell nuclei were stained with DAPI. The eye diameter (B), the eye area (C), the retina area (D), the optic nerve area (E), and the optic tract area (F) were measured and compared between control and Sert-Cre cDKOs and showed no significant differences. (G) Retina sections from mouse P0 control or Sert-Cre cDKO were stained for Zic2. (H) The number of Zic2^+^ cells per section was not different between genotypes. (I) Flat mount preparation of adult control or Sert-Cre cDKO retinas showing tdTomato^+^ cells. (J) The tdTomato^+^ cell density shows no difference between control and Sert-Cre cDKO. Data are mean±s.e.m and dots represent individual animals. Mann-Whitney tests were performed in B, C, D, H and J. Two-way ANOVA was performed in E and F. Scale bars: 250 μm in A; 50 μm in G; 100 μm in I.

Since early deletion of Numb/Nbl in retinal progenitors showed that Numb/Nbl are required for the normal production of retinal cell types (Kechad et al., 2012), we next examined whether iRGC specification and survival are affected in Sert-Cre cDKO. For this, we stained P0 retinal sections from control or Sert-Cre cDKO for Zic2, a transcription factor that specifies the ipsilateral fate of RGCs (Herrera et al., 2003). Quantification of the number of Zic2^+^ cells showed no changes between controls and Sert-Cre cDKO (Figs. 2G, H). We also looked in adults (3-4 months old) and observed that the number of Sert-Cre tdTomato^+^ cells was not different between controls and Sert-Cre cDKO (Figure 2I, J), demonstrating that *Sert-cre* mediated Nb/Nbl inactivation does not affect the production or survival of iRGCs in the retina. The difference in the phenotype of Numb/Nbl inactivation between the study by Kechad et al. and our study is due to the difference in onset of expression and cell-type specificity of the two distinct Cre: while *Sert-Cre* starts to be expressed at E14.5 in committed iRGCs (Koch et al., 2013; Peng et al., 2018), the *alpha-Pax6-Cre* starts to be expressed at E9 in early retinal progenitors (Kammandel et al., 1999).

### Numb/Nbl is required for ipsilateral axon segregation *in vivo* and Shh-induced iRGCs growth cone collapse

iRGC axons are repelled at the level of the optic chiasm and thus ultimately project ipsilaterally to the thalamus. One of the guidance cues known to control this repulsion is Shh (Fabre et al., 2010). Shh is produced by cRGCs and transported along their axons to the optic chiasm where it repels iRGC axons locally (Peng et al., 2018). This effect of Shh is mediated by its receptor Boc (Fabre et al., 2010). In the developing spinal cord, Shh-mediated chemoattraction of commissural axons also acts via Boc and requires Numb as an endocytic adaptor (Ferent et al., 2019). However, the molecular mechanism underlying the repulsion induced by Shh via Boc in iRGC growth cones remains unknown. Therefore, we wondered whether Numb/Nbl might also play a role in the repulsion of iRGC axons at the optic chiasm. To investigate the projections of ipsilateral axons from the eye to the brain, we injected DiI into one eye and quantified DiI fluorescence intensities in contralateral and ipsilateral optic tract sections at P0. Compared to the control, Sert-Cre cDKO resulted in a decrease in the ipsilateral fraction (control: 14.67% ± 0.67%, Sert-Cre cDKO: 10.94% ± 0.42%; n≥4 animals; p = 0.0141) (Fig. 3A, B). To further characterize this decrease in the segregation of RGC axons at the chiasm, we examined the distribution of terminal projections in the dorsal lateral geniculate nucleus (dLGN) of the thalamus by whole-eye anterograde labeling using cholera toxin subunit B (CTB) coupled to fluorophores. Since each eye is injected with a CTB conjugated to a different fluorophore, retinal axons can be labeled in an eye-specific manner and their projections analyzed in the dLGN (Figure 3C). Ipsilateral axons are shown in green and contralateral axons are shown in red. The fluorescence intensity of the ipsilateral projections was decreased in the dLGN of Sert-Cre cDKO compared to the control. This was confirmed by analyzing the percentage of the dLGN territory occupied by ipsilateral inputs (Figure 3D). We performed this analysis using different fluorescence threshold and observed a consistent trend. When the ipsilateral detection threshold was set to 35, the percentage of dLGN area occupied by ipsilateral terminals showed a 36% decrease in Sert-Cre cDKO compared to controls (p=0.0367, Figure 3E). Importantly, the total dLGN area remained unchanged (Figure 3F).

**Figure 3:**
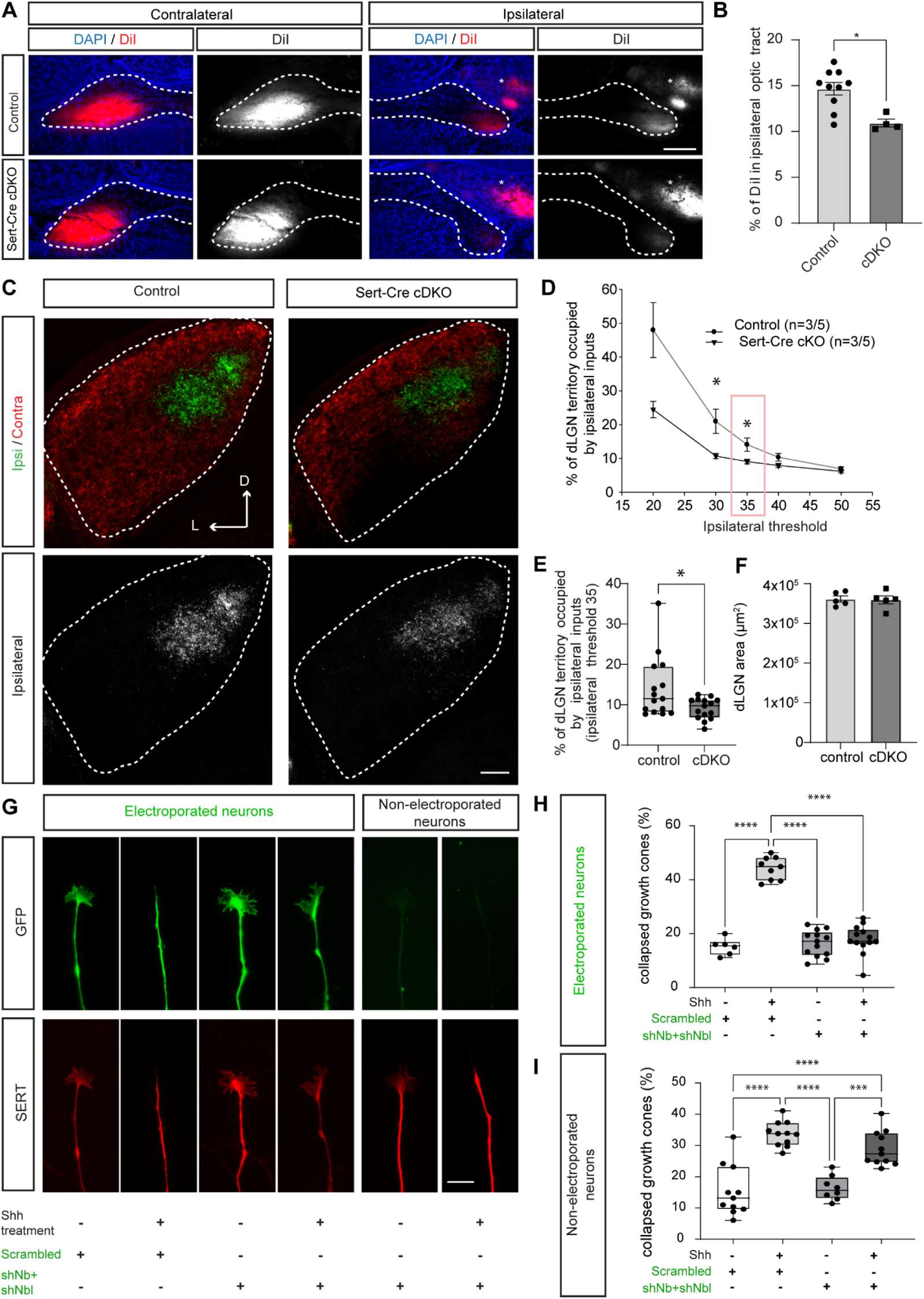
Numb/Nbl is required for ipsilateral axon segregation *in vivo* and Shh-induced iRGCs growth cone collapse. (A) Optic tract sections from P0 control and Sert-Cre cDKO animals injected with DiI. (B) Decreased percentage of DiI fluorescence in Sert-Cre cDKO ipsilateral optic tract compared to controls (mean±s.e.m). (C) Retinogeniculate projections in control and Sert-Cre cDKO dLGN at P0, with ipsilateral projections in green and contralateral projections in red. Binary images of ipsilateral projections are shown below. (D) Percentage of ipsilateral segregated inputs in the dLGN as a function of threshold (mean±s.e.m). (E) At a threshold of 35, Sert-Cre cDKO mice show a decreased percentage of dLGN territory occupied by ipsilateral projections. * Mann-Whitney test p=0.0367. (F) No difference in dLGN area between control and Sert-Cre cDKO (mean±s.e.m). (G) iRGCs from ventro-temporal retina explants electroporated with scrambled or shNb and shNbl (green), identified by SERT immunostainings (red). (H) Quantification of the percentage of electroporated iRGC collapse after Shh or control stimulation. One-way ANOVA, dots represent individual coverslips. (I) Quantification of the percentage of non-electroporated iRGC collapse after Shh or control stimulation. Scale bars: 200 μm in A; 100 μm in C; 20μm in G.

To determine whether Numb is required for Shh-dependent repulsion in the context of iRGC axon guidance, we performed Shh-induced collapse assays on retinal explants. We knocked-down Numb and Nbl using specific shRNAs (which we validated in a previous study (Ferent et al., 2019)) by electroporating the shRNA expression constructs into ex-utero E14.5 retina. Retinas were dissected and explants were grouped according to the orientation of the eye: ventro-temporal (VT) explants containing iRGCs were separated from non-VT explants containing all other regions. Immunostaining for SERT indicated the ipsilateral identity of the electroporated axons, which themselves were identified by GFP expression (GFP is expressed by the shRNA plasmid). The specificity of these staining was further validated by co-staining with an anti-Shh, which labels cRGCs and shows no colocalization with SERT (Figure S1A and B). Similar to what we showed previously, a 30-minute stimulation with Shh induced a significant increase in the proportion of collapsed growth cones from iRGCs electroporated with a scrambled control plasmid (Fabre et al., 2010). However, this effect was blocked when Numb/Nbl were knocked-down (Figure 3G, H). We also analyzed non-electroporated growth cones from the same explants and confirmed that GFP-negative iRGCs retained their sensitivity to Shh, as the percentage of collapse of these growth cones increased in response to Shh (Figure 3G, I).

Together, these results demonstrate that Numb/Nbl are required for proper ipsilateral retinal projection segregation at the optic chiasm and for Shh-mediated collapse of iRGCs.

### Numb/Nbl is required for post-midline crossing guidance in the developing spinal cord

In addition to the developing visual system, Shh also has a repulsive function in the developing spinal cord. After commissural axons reach the midline, they switch their sensitivity to Shh from attraction to repulsion (Yam et al., 2012). Shh then acts as a repellent to make commissural axons turn anteriorly.

Since Numb/Nbl is required for commissural axons to be initially attracted by Shh (Ferent et al., 2019), and that Numb/Nbl is also required for Shh-mediated repulsion in the visual system (Figure 3), we wondered whether Numb/Nbl is also required for repulsion of commissural axons after midline crossing. We first examined the expression of Numb and Nbl in commissural neurons. Using commissural neurons isolated from E13.5 rat spinal cords and cultured for four days (4DIV), we performed immunostaining for Numb/Nbl and co-staining for filamentous actin (phalloidin). At 4DIV, commissural axons are repelled by Shh (Yam et al., 2012). We showed that Numb/Nbl is present at the growth cone level in vesicle-like structures, in filopodia and in the axonal shaft (Figure 4A). RNA sequencing (RNA-seq) revealed significant expression of Numb and Nbl in commissural neurons, similar to the levels of Ptch1 (Figure 3B), both at 2DIV, when axons are attracted to Shh, and at 4DIV, when they are repelled by Shh. Using mouse embryos at E11.5, we also stained for Numb/Nbl in vivo and for L1, a marker of post-crossing commissural axons (Figure 4C). We observed that L1 staining colocalizes with the Numb/Nbl signal. Taken together, this shows that Numb/Nbl is highly expressed by commissural neurons after their axons have crossed the midline. Next, we investigated if Numb and Nbl are required for guidance after midline crossing. Because Numb is required for commissural axon guidance to the midline (Ferent et al., 2019), we analyzed conditional mutant embryos in which Numb/Nbl is removed from dI1 commissural neurons by crossing *Nb; Nbl* floxed mice (Wilson et al., 2007) with mice expressing the Math1-Cre driver (Matei et al., 2005). First, the fluorescence intensity of L1 staining in Math1-Cre cDKOs did not change compared to controls (Figure 4D, E), nor did the thickness of the ventral commissures as measured by TAG1 immunostaining (Figure 4F, G). As we have previously shown with a ROBO3 staining (Ferent et al., 2019), this indicates that although the inactivation of Nb/Nbl induces guidance errors on the way to the midline, the majority of axons still reach the floor plate. To assess the role of Numb/Nbl in antero-posterior guidance, we labeled commissural axons in E12.5 embryos with DiI. This technique visualizes the trajectory of commissural axons after crossing. We observed a significant decrease in the ratio of anterior-guided axons to the total fluorescence of both anterior- and posterior-guided axons in the Math1-Cre cDKO compared to the controls (Figure 4H, I). This indicates an increase in misrouted axons that turn toward the posterior part of the neural tube in absence of Nb/Nbl. Thus, our results reveal that Numb/Nbl play a role in the guidance of post-crossing commissural axons along the longitudinal axis.

**Figure 4:**
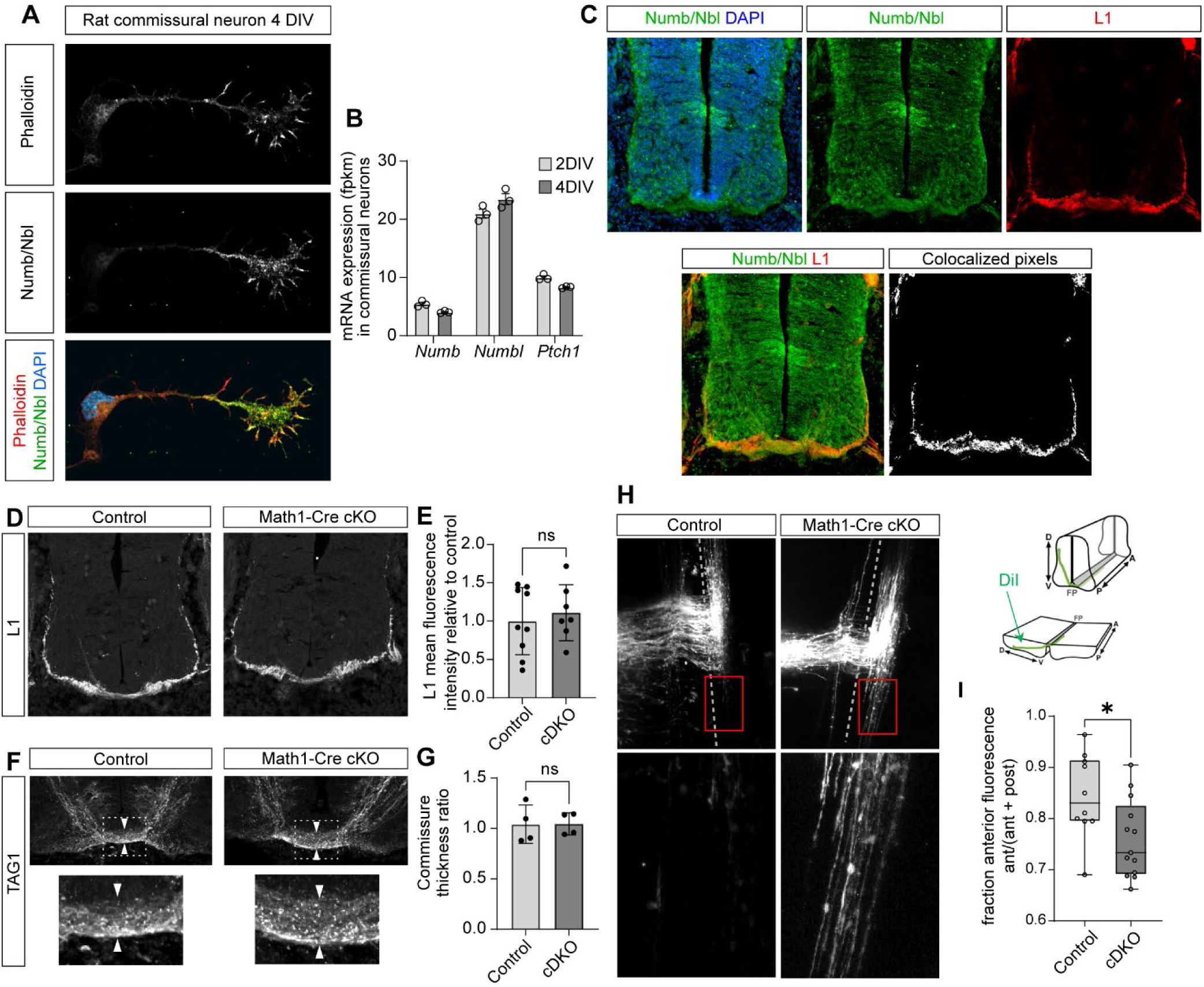
Numb/Nbl is required for post-midline crossing guidance in the developing spinal cord. (A) Rat commissural neurons cultured *in vitro* for 4 days (4 DIV) were immunostained for Numb/Nbl and stained with phalloidin for actin visualization. Nuclei were detected by DAPI. (B) Mean mRNA expression (fpkm ± SEM) of Numb, Nbl, and Ptch1 in dissociated commissural neurons at 2 and 4 DIV (n = 3 independent experiments). (C) Mouse e11.5 neural tube sections immunostained for Numb/Nbl and L1 to label commissural axons after crossing. (D) L1 staining of E11.5 spinal cord sections from control and Math1-Cre cDKO. (E) Mean L1 fluorescence intensity relative to control shows no difference in Math1-Cre cDKO mice compared to controls (Mann-Whitney, dots represent individuals). (F) TAG1 staining of E11.5 spinal cord sections from Math1-Cre cDKO and control show ventral commissure morphology. (G) Commissure thickness is unchanged in Math1-Cre cDKO mice compared to controls (Mann-Whitney, dots indicate individuals). (H) DiI labeling of post-crossing commissural axons in open book preparations of control and Math1-Cre cDKO mouse embryos, with magnification of the boxed region. Red boxes indicate magnified areas, vertical dashed lines indicate the midline. Anterior is up. (I) Relative fluorescence of anterior-directed axons versus total fluorescence of anterior- and posterior-directed axons (n ≥ 10 embryos per genotype), * Mann-Whitney test p=0.0214, dots represent individuals.

## CONCLUSION

Collectively, our data indicate that Numb/Nbl are required for repulsive axon guidance in two distinct neurodevelopmental systems: iRGC segregation in the visual system and antero-posterior guidance after midline crossing in the spinal cord. In both of these paradigms, Shh plays an important role in inducing growth cone repulsion. We also show that Numb/Nbl also play a critical role in Shh-mediated iRGC collapse, directly linking its function in repulsion and the Shh signaling pathway. In addition to the role of Numb/Nbl in Shh-mediated attraction (Ferent et al., 2019) and canonical pathway activation (Liu et al., 2024), we now demonstrate that Shh-mediated axon repulsion also relies on Numb/Nbl, positioning Numb/Nbl as an essential component of the Shh signaling pathway in multiple neurodevelopmental functions.

## MATERIALS AND METHODS

### Animals

All animal work was performed in accordance with the Canadian Council on Animal Care Guidelines and approved by the IRCM Animal Care Committee. Staged pregnant Sprague Dawley rats were obtained from Charles River (St. Constant, Canada) and staged pregnant wild type mice used for explants were obtained from Janvier labs (France). Transgenic mice were maintained in the IRCM specific pathogen-free animal facility. All mouse lines have been previously described: Math1-Cre (Matei et al., 2005), Slc6a4 (ET33-SERT)-Cre (Gong, Doughty et al. 2007, obtained from the Mutant Mouse Regional Resource Center (MMRRC) of UC Davis, Numb conditional allele and NumbLike conditional allele (Wilson et al., 2007, obtained from The Jackson Laboratory), ROSA26-tdTomato (Madisen, Zwingman et al. 2010, kindly provided by Dr. Ying Zhang at the Dalhousie University). When ROSA26-tdTomato was used, control genotype is Sert-cre; Nb lox/+; Nbllox/lox. Embryonic day 0 (E0) was defined as midnight of the night before a plug was found.

### Immunostaining

P0 eyes sections were stained with anti-Zic2 (a kind gift from C. Mason). Embryo sections were immunostained with anti-Numb/NumbLike antibody (Abcam, ab14140), anti-Tag-1 (R&D Systems, AF4439) and anti-L1 (Chemicon MAB5272). Retinal explants were permeabilized and blocked with 0.25% (v/v) Triton and 3% (w/v) BSA in PBS, then immunostained with antibodies against: GFP (Aves Labs, GFP1020) followed by a secondary antibody coupled to AlexaFluor 488 (Jackson ImmunoResearch), SERT (Santa-Cruz, sc-1458) followed by a secondary antibody coupled to AlexaFluor 594 (Abcam), and βIII-tubulin (Abcam, ab18207) followed by a secondary antibody coupled to Cy5 (Jackson ImmunoResearch). Antibodies were diluted in PBS supplemented with 0.1% (v/v) Triton and 1% (w/v) BSA. Dissociated rat commissural neurons were immunostained with anti-Numb/NumbLike antibody (Abcam, ab14140). Filamentous actin was detected with phalloidin (Sigma, P1951).

### DiI tracing

For iRGC projection measurement in the optic tract, mono-ocular DiI crystals-filling of the retina was performed at P0 on fixed tissue, as previously described (Fabre et al., 2010; Plump et al., 2002). After allowing the dye to diffuse, 30 µm-thick coronal sections just caudal to the optic chiasm were performed and the amount of DiI fluorescence signal was quantified in the contralateral and ipsilateral optic tracts.

All images were acquired under conditions where the pixels were not saturated using a Leica DM4000 microscope (Leica Microsystems GmbH, Wetzlar, Germany) and an Orca ER CCD camera (Hamamatsu Photonics, Hamamatsu City, Japan). Fluorescence measurements were taken blind to the genotype on ten consecutive sections per animal using Volocity version 4.3 (Improvision, Waltham, MA, USA). The ipsilateral index was calculated by dividing the fluorescent intensity in the ipsilateral optic tract by the sum of the fluorescent intensity in ipsilateral + contralateral optic tracts.

For the post-crossing guidance assay in the spinal cord, the neural tubes were dissected from the embryo and fixed at least overnight at 4°C in 4% paraformaldehyde in PBS. After fixation, a small amount of 1,1’-dioctadecyl-3,3,3’,3’-tetramethylindocarbocyanine perchlorate (DiI, Molecular Probes, Eugene, OR) dissolved in ethanol (10mg/ml) was inserted to the medial neural tube dorsal of the motor column to label several individual cohorts per embryo (around 5-9) at multiple levels along the AP axis (Farmer et al., 2008). The DiI was allowed to diffuse for 2 days at 4°C. After diffusion of the dye, the neural tubes were mounted in an open-book configuration and imaged on a confocal microscope (Leica LSM700). Analysis of axon guidance and image processing was performed on the resulting Z-stacks in ImageJ (NIH). All images shown are maximum projections of the Z-stacks.

### CTB tracing

Anterograde CTB labeling were performed as previously described (Jaubert-Miazza et al., 2005; Rebsam et al., 2009). Briefly, adult mice were deeply anesthetized with a mixture of isoflurane (5% for induction and 2% for maintenance) and a 1:1 flow ratio of air/O2 (1l/min). Eyes were injected with a glass pipette intravitreally with 5µl of 0.2% cholera toxin B subunit (CTB) conjugated to Alexa Fluor 594 or 488 (Invitrogen by Life Technology, Carlsbad, CA, USA) diluted in 1% DMSO. After 3 days, mice were anesthetized and perfused with 4% PFA in 0.1 M PB before dissection of brain and retinas. Brains were post-fixed overnight, embedded in 4% agarose, and sectioned (80 µm) with a Leica VT1000S vibratome. Coronal sections were directly mounted in Mowiol and imaged on a confocal microscope (Leica LSM700). As Alexa Fluor 594-labeled contralateral/Alexa Fluor 488-labeled ipsilateral projections had a better signal-to-noise ratio, the analyses were conducted on these images, as already described (Muir-Robinson et al., 2002; Rebsam et al., 2009; Torborg and Feller, 2004). Quantification was performed on Z-stack images from three consecutive coronal sections through the region of the dLGN containing the greatest extent of the ipsilateral projections. For this, the boundary of the dLGN was delineated in order to exclude label from the intrageniculate leaflet, the ventral lateral geniculate nucleus, and the optic tract. Using ImageJ software (NIH), the proportion of dLGN occupied by ipsilateral axons was quantified as a ratio of ipsilateral pixels to the total number of pixels in the dLGN region using a multi-threshold analysis.

### Retinal explant cultures and collapse assay

Retinas of E14.5 mouse embryos were electroporated with Numb and Numb like or scramble shRNA (1 µg·µl^-1^ for each plasmid) using 5 pulses of 45 V during 50 ms every 950 ms with an ECM 830 square wave electroporator (BTX). shNb and shNbl have been previously validated in our hands (Ferent et al, 2019). Retinas were dissected and kept 24 hours in culture medium (DMEM-F12 supplemented with 1 mM glutamine (Sigma Aldrich), 1% penicillin/streptomycin/AmphotericinB (Cytiva), 0.001% BSA (Sigma Aldrich) and 0.07% glucose), in a humidified incubator at 37°C and 5% CO2. The following day, the retinas were cut into 200 µm squares with a razorblade and explants were plated on glass coverslips coated with 100 µg·ml^-1^ poly-lysine and 20 µg·ml^-1^ laminin (Sigma Aldrich). Cells were cultured for 24 hours in culture medium supplemented with 0.5% (w/v) methylcellulose and B-27 (1/50, Life technologies). Retinal explants were treated with 240 ng·ml^-1^ SHH C24II (R&D Systems, 1845-SH) diluted in warm culture medium for 30 minutes before fixation with 4% (w/v) PFA in PBS for 30 minutes.

### Commissural neuron cultures

Spinal commissural neurons were dissected and cultured as described previously (Langlois et al., 2010).

### Quantification and statistical analysis

Statistical analyses were performed with GraphPad Prism 10 (La Jolla, CA). Student’s t-test or Mann Whitney’s test were used when there were two groups in the dataset. To compare multiple groups in a dataset, one-way ANOVA was used. The statistical analysis used in each experiment and the definition of n are stated in the figure legends.

## ACKNOWLEDGMENTS

We acknowledge the Imaging Platform of Institut du Fer à Moulin for their help in fluorescence imaging. We thank Drs. Carol Mason for providing us with the Zic2 antibody. The 5E1 antibody was obtained from the Developmental Studies Hybridoma Bank developed under the auspices of the NICHD and maintained by the University of Iowa.

## AUTHOR CONTRIBUTIONS

Conceptualization: T.D, J.F.; Methodology: T.D, S.B., J.F.; Formal analysis: T.D, S.B., J.F.; Investigation: T.D, S.B., J.F.; Writing - original draft: J.F, F.C.; Supervision: J.F, F.C.; Funding acquisition: J.F, F.C.

---

**Figure S1:**
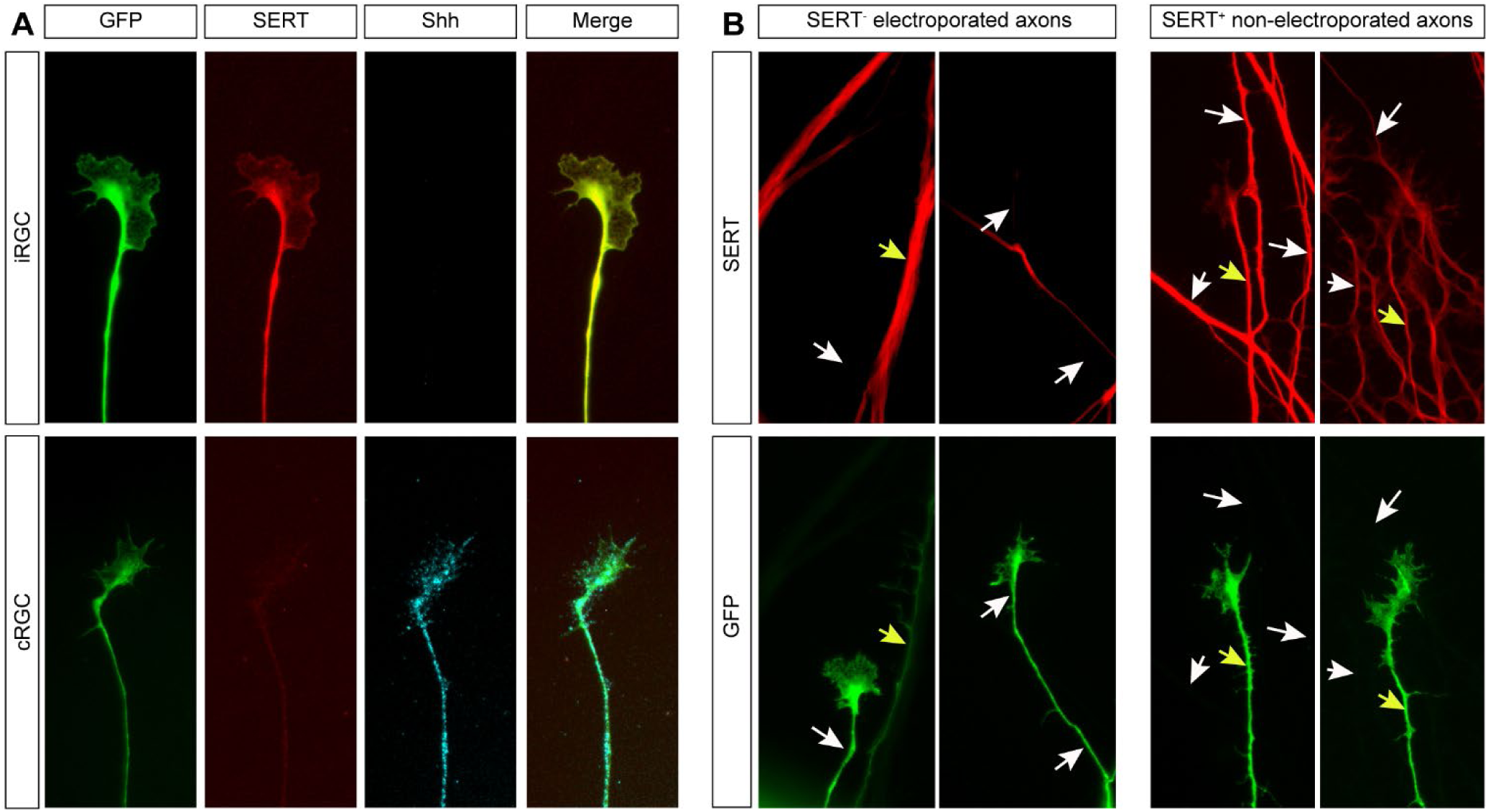
Identification of iRGCs from ventro-temporal retinal explants in vitro. (A) Immunostaining for SERT (red) and Shh (blue) of GFP-electroporated (green) axons of RGCs grown from retinal explants at E14.5 + 2 days *in vitro*, either from the ventro-temporal region (iRGC) or the dorso-medial region (cRGC). (B) Examples of immunostaining for SERT (red) and GFP-electroporated (green) axons of RGCs grown from retinal explants at E14.5 + 2 days *in vitro,* showing double labeled axons, thus electroporated iRGCs (GFP^+^, SERT^+^, yellow arrows), electroporated cRGCs (GFP^+^, SERT^-^, left panels, white arrows) and non-electroporated iRGCs (GFP^-^, SERT^+^, right panels, white arrows).

